# HIP1 mediates oncogenic transformation and cancer progression through STAT3 signalling

**DOI:** 10.1101/2020.07.09.191734

**Authors:** Roheet Bantval Rao, Antonio Ramos-Montoya, Helen Scott, Lorraine Berry, Stefanie Reichelt, Stewart MacArthur, Roslin Russell, David E Neal, Emma Evergren, Ian G Mills

## Abstract

**Background:** Huntingtin-interacting protein 1 (HIP1) is an adaptor protein involved in transcriptional regulation and receptor-mediated endocytosis. Overexpression of HIP1 transforms cell and is associated with, increasing grades of prostate cancer (CaP) and poor patient outcomes. However, the precise mechanism for the role of HIP1 in prostate cancer progression remains unknown.

**Methods:** Using a phospho-kinase antibody array we identified changes in signalling associated with HIP1 overexpression PNT1 cells. For validation Western blots were used together with knockdown or inhibitor treatments and phenotypic assays for cellular transformation. The cell line was xenografted to assess tumour growth. Gene expression microarray analysis of the cell line was used to identify perturbations in transcript levels.

**Results:** Here we demonstrate cellular transformation and phenotypic effects of HIP1 overexpression in a benign prostate epithelial cell line to be dependent on STAT3 signalling. *In vivo* xenografts confirmed the cellular transformation phenotype. Gene expression analysis revealed serum protein GDF15 to be a marker of prostate cancer tumorigenesis in our model. We present a HIP1-STAT3-GDF15 axis in our pre-clinical model that mediates cellular transformation and tumorigenesis.

**Conclusion:** Our findings provide a model defining the functional effects of increased HIP1 expression in prostate tumorigenesis and progression. This model implicates increased STAT3 signalling in HIP1-dependent prostate carcinogenesis and identifies GDF15 as a secreted factor supporting this process. The role of HIP1-STAT3-GDF15 signalling may extend to other epithelial cancers shown to overexpress HIP1; such as gliomas, colon and breast cancer where STAT3 is an emerging oncology drug target.

## Background

Huntingtin-interacting protein 1 (HIP1), a highly conserved 116kDa protein, is an adaptor protein principally involved in clathrin-mediated internalisation of cell surface receptors facilitated by its interactions with various components of the endocytic machinery (1–5). HIP1 overexpression is associated with a variety of epithelial malignancies such as breast, colon, prostate cancer and gliomas (6–8). In prostate cancer, HIP1 expression correlates with cancer progression and poorer outcomes (6, 9). Furthermore, HIP1 autoantibodies are detected with higher frequency in patients with prostate cancer (CaP) compared to cancer-free individuals suggesting its possible use as a serum biomarker for the detection of primary CaP (10, 11). HIP1 overexpression in mouse models of prostate cancer drives tumourigenesis and progression of these tumours (10). Together these data demonstrate an important role for HIP1 in prostate cancer, which may be due to aberrant growth factor endocytosis and signalling as shown in fibroblasts and prostate epithelial cells where overexpression of HIP1 results in sustained signalling (12) (9). We have previously shown an additional mechanism that does not involve HIP1’s canonical function in endocytosis, but requires a nuclear translocation of HIP1 where it associates with the androgen receptor (AR) upon androgen stimulation (8).

In this study we demonstrate that HIP1 overexpression results in increased STAT3 phosphorylation and its nuclear translocation, which is mediated in part by downstream FGF signalling. We also demonstrate that STAT3 activity is critical for the phenotypic effects of HIP1 overexpression, and identify increased GDF15 expression upon cellular transformation.

## Methods

### Reagents and Antibodies

The following gene expression constructs were used: GFP-HIP1 plasmid previously described (8) and Flag-HIP1 constructs previously described (9).

The following antibodies were used: GFP (rabbit polyclonal, AbCam #ab6556), b-actin (mouse monoclonal AC-15, AbCam #ab6276-100), HIP1 (mouse 4B10, Santa Cruz # sc-47754), STAT3 (rabbit, Cell signalling technologies #9132), p-STAT3 (T705) (rabbit, Cell signalling technologies #9131S), Phospho-FGF R (Y653/Y654) (Affinity Purified Polyclonal Rabbit IgG, R&D systems #AF3285), phospho-JAK2 (Y1007 + Y1008) (rabbit E132, Abcam #ab32101). For the GDF15 conditioned media experiment, FGFR4 (FR4Ex) antibody (rabbit, Santa Cruz #sc-73995) was used. The secondary antibodies used for western blotting were horseradish peroxidase (HRP)-conjugated antibody against mouse immunoglobulin G (IgG) and (HRP)-conjugated antibody against rabbit IgG from Dako.

The following primers were used for expression qRT-PCR: GAPDH (5’-GAAGGTGAAGGTCGGAGTC-3’, 5’-AAGATGGTGATGGGATTTC-3’), GDF15 (5’- TGTCGCTCCAGACCTATGATGA-3’, 5’-AATCGGGTGTCTCAGGAACCTT-3’), HIP1 (5’- TGCTCTGCTGGAAGTTCTG-3’, 5’-CTGGCGGTCACTCATCTG-3’).

### Generation of Stable Cell Lines and RNA interference knockdown

PNT1a, DU145 and LNCaP cell-lines were grown in RPMI (Gibco #21875-034) supplemented with 10% foetal bovine serum (Gibco #10270-106); 0.8mg/ml G418 was supplemented for maintaining PNT1a-HIP1 and PNT1a-EV cell lines while 0.8mg/ml G418 and 2.0μg/ml of Puromycin (Sigma # P9620) was supplemented for maintaining the PNT1a stable knockdowns of HIP1. Stable overexpression of HIP1 was generated by transfection using Nucleofector technology (Lonza). Stable knockdown of HIP1 was generated by nucleofection (Lonza) of the two shHIP1 constructs (Origene, HuSH pRS plasmids #TR312457) in PNT1A and DU145 followed by selection with 2.0μg/ml final concentration of Puromycin (Sigma #P9620) added to the culture medium. Following selection, multiple clones within the same transfection dish were allowed to proliferate and two such multiclonal PNT1A-shHIP1 lines were established. Stable PNT1AshHIP1 knock-down cell-lines were maintained in RPMI supplemented with 10% FCS, 0.8mg/mL Geneticin G418 sulphate (Gibco #10131-019), and 1.0 μg/mL concentration of Puromycin (Sigma #P9620) selection antibiotic. Similarly, PNT1A-HIP1 cells transfected with plasmid expressing scrambled hairpin RNA (Origene, HuSH pRS plasmids #TR312457) supplied within the same kit, and selected as above served as control. Multiclonal lines, expressing two different shHIP1 hairpins were generated to prevent clonal effects. The knock-downs were confirmed by determining HIP1 mRNA expression by real-time PCR. DU145 shHIP1 knock-down cell lines were maintained in RPMI supplemented with 10% FCS, 1.75 μg/mL concentration of Puromycin (Sigma #P9620) selection antibiotic. Pooled human HIP1 siRNA (Santa Cruz #sc-41982); pooled STAT3 siRNA (Qiagen, #FlexiTube GeneSolution GS6774); scrambled siRNA (Dharmacon; ON-TARGETplus non-targeting siRNA#2) were used for transient RNA interference experiments and transfected using Nucleofector technology (Lonza).

### Proteome array

The Prosphoproteome array was used according to the manufacturer’s instructions (R&D systems, # ARY003) with 400μg of protein used per membrane. Membranes were scanned with Labscan™ and image analysis was done with ImageQuant™ software.

### Western blotting

Total protein lysates were prepared by lysing cells on ice in 1% NP-40 lysis buffer (50mMTris, pH 6.8, 150mM NaCl, 1mM NaVO4, 10mM DTT (Dithiothreitol), 1mM PMSF (Phenylmethanesulfonyl fluoride), 1 tab Phos-stop EDTA free(Roche#4906837001), and complete protease inhibitor (Roche) per 50mL, 1% NP-40). Protein concentrations in supernatant was quantified using Coomassie reagent (PierceTM, #23200) and interpolated from a standard curve obtained using bovine serum albumin. Total cell lysate was resolved by SDS-PAGE using 4-12% gels; 8% gels (PierceTM),transferred to nitrocellulose or PVDF membranes (Invitrogen) using the I-Blot Dry Transfer System (Invitrogen). ECL plus reagent (GE Healthcare) was used to visualise Western blots. Quantification of the bands was performed bydensitometry with Image-J software (http://rsbweb.nih.gov/ij/); actin bands were used for normalisation to ensure equal protein loading in all lanes.

### Colony Formation Assay

CytoSelect™ 96-Well Cell Transformation Assay (Fluorometric Assay), Cell Biolabs Inc. was used for assessing anchorage independent growth/soft agar colony formation using the manufacturer’s instructions. Briefly, cell lines were trypsinized and resuspended in media, counted in the Vi-cell^TM^, diluted to achieve identical concentrations of each cell line and the relative fluorescence units (RFU) of each cell suspension were measured as per the manufacturer’s instructions. Samples were analysed in triplicate on Day 7, to assess the soft agar colony formation for each cell line. The ratio of the RFU between the control and test cell lines at seeding was used to correct the final RFU measured in the assay to adjust for any variation at the point of cell seeding.

### Migration Assay

The CytoSelect™ 96-Well Cell Migration and Invasion Assay (8μm, Fluorometric Format, Cell Biolabs Inc.) was used for assessing cell migration. Cell lines were serum starved for 24 hours, trypsinized, and resuspended in serum free media prior to being counted in the Vi-cell^TM^. The cell suspensions were diluted to achieve identical concentrations of each cell line. Standard does curves for each cell line was plotted as per the manufacturers’ instructions. The fluorescence (RFU) was measured after 24 hours to assess cell migration towards media supplemented with foetal calf serum, for each cell line in triplicate. The ratio of the fluorescence between the control and test cell lines at seeding was used to correct the final measurement in the assay to adjust for any variation at the point of seeding.

### MTS assay for drug IC50

CellTiter 96^®^ Aqueous Non-Radioactive Cell Proliferation Assay (MTS) (Promega, # G5421) was used. Cells were seeded in 96 well plates at the required density following cell counting. Manufacturer’s instructions were followed. Absorbance was read at 490nm after 1 hour incubation.

### Microarray analysis

Total RNA from PNT1a stable cell lines and LNCaP cell lines transiently transfected with HIP1 three days post-transfection was extracted using Qiagen miRNeasy kit according to the manufacturer’s instructions. Quality control was performed with an Agilent Bioanalyser. cDNA was generated and biotin labelled using the Illumina TotalPrep RNA Amplification Kit, according to the manufacturer’s instructions. Hybridization and scanning were performed using Standard Illumina protocols. Expression analysis was carried out on the Illumina humanHT-12 v3 beadchip. Quality control and processing was carried out in R using the Bioconductor package beadarray. Differential expression analysis was carried out with the Bioconductor package limma. Network analysis was done using proprietary software (curated) GENEGO Metacore™.

### Quantitative Real-Time PCR

All RNA extractions were done using the miRNeasy kit (Qiagen) as per manufacturer’s protocols. Quantitative real-time PCR was performed following cDNA preparation using ABI Mastermix as per manufacturer’s protocols followed by SYBRgreen chemistry (Applied Biosystems, 2x SYBRgreen master mix) in an ABI7900 instrument (Applied Biosystems).

### Immunoflorescence and Laser scanning cytometry

Cells were seeded in equal numbers in Ibidi ibiTreat™ 8 well μ-slides (Ibidi GmbH, Germany), fixed in 4% para-formaldehyde (formalin-free) for 10min at room temperature, methanol permeablised and 7pprox-stained with anti-STAT3 antibodies (1:100) overnight followed by incubation with secondary Alexa488-tagged antibodies and Hoechst staining (1:5000 in PBS). Quantitative imaging cytometry using an iCys Research Imaging Cytometer (CompuCyte, Cambridge, MA) with iNovator software (CompuCyte) was used to collect images using a 60x objective (Uplan FLN 0.90NA, Olympus, Tokyo, Japan) with 1.0 μm step size, giving a field size of 1000μm x 132.1μm^2^. A minimum of 162 fields were scanned. Nuclei were contoured using a binary threshold in the 405nm (Hoechst) channel according to size (40 to 800μm^2^) and intensity. A water shed filter (0.1; 3-8 pixels) was incorporated within the protocol to separate touching nuclei. The amount of Alexa Fluor 488 staining was measured within the nuclear contour.

### Conditioned media and GDF15 ELISA

Cells were cultured in OptiMEM media for 24 hours and conditioned media collected, cleared and concentrated to 1 ml in a VIVASPIN™ 20 column (GE Healthcare). The concentration of GDF15 was measured using an ELISA assay for Human GDF-15 DuoSet (R&D systems, # DY957) as per the manufacturer’s instructions on a Tecan INFINITE^®^ 200 Pro spectrophotometer. A standard curve was generated for each plate. Protein quantification and analysis was done in Graphpad PRISM using a four-parameter sigmoidal (logistic) model and interpolation from the standard curve.

### Mice, Xenografts, and Imaging

Mice Immunocompromised NOD-SCID-GAMMA (NSG) male mice (Charles River, Wilmington, MA) were used for tumour implantation. Mice were maintained in the Cancer Research UK Cambridge Institute Animal Facility. All experiments were performed in accordance with national guidelines and regulations, and with the approval of the animal care and use committee at the institution. All animal procedures were carried out in accordance with University of Cambridge and Cancer Research UK guidelines under UK Home Office project license 80/2301. PNT1a-EV and PNT1a-HIP1 cells were stably transfected with a transposon for expression of luciferase as described previously (Mills et al., 2005). Cells were selected and maintained in RPMI supplemented with 0.8mg/ml G418 and 2μg/ml Puromycin. Eleven-week-old mice were xenografted with 1.7 million cells of PNT1a-EV (n=10) or PNT1a-HIP1 (n-20). Bioluminescent imaging of the mice was done using the Xenogen as per manufacturer’s instructions. Mice were injected with Luciferin 30 minutes prior to imaging. Imaging parameters used for all the groups was identical and carried out fortnightly. For the assessment of GDF15 in mice sera, approximately 100μl of mouse blood was collected at 26 weeks post-xenograft and serum obtained as previously described (13).

### Statistical analysis

Statistical calculations were performed with Graphpad PRISM software. The differences between groups were calculated using Student’s t-test or ANOVA.

## Results

### HIP1 overexpression enhances STAT3 phosphorylation

HIP1 overexpression in a benign immortalised prostate epithelial cell line (PNT1A) has been shown to enhance FGFR4 stabilisation and was implicated in the phenotypic effects of HIP1 overexpression (9). Using a phospho-kinase antibody array, which detects relative protein phosphorylation levels of 46 sites in kinases and substrates, we investigated the downstream effects of FGFR stabilisation caused by stable HIP1 overexpression in PNT1A cell lines (PNT1A-HIP1) cultured in full serum compared to control cells (PNT1A-EV). We saw a >1.5 fold increase in tyrosine phosphorylation (Y705) of STAT3, serine phosphorylation (S473) in Akt, threonine phosphorylation (T174) in AMPK1α, and tyrosine phosphorylation of PLC-γ1 **(Figure 1A and Supplementary Figure S1A)** in PNT1A-HIP1 cells. Western blots validated the increased STAT3 phosphorylation (Y705) levels arising from HIP1 overexpression **(Figure 1B**). Transient overexpression of HIP1 in the PNT1A-EV cell line also led to increased STAT3 phosphorylation (**Figure 1C**). Since HIP1 has previously been shown to promote cellular transformation and tumorigenesis when overexpressed in the NIH-3T3 cell-line we also transiently transfected HIP1 into this line. HIP1 overexpression in the NIH-3T3 line led to an increase in phosho-STAT3 levels (**Figure 1D**). Stable knockdown of HIP1 in the PNT1A-HIP1 cells reduced STAT3 phosphorylation (**Figure 1E**). Together these data highlighted that increases in STAT3 phosphorylation were due to HIP1. Of the two stable HIP1 knockdown clones that were generated, clone 1 showed the more significant reduction in HIP1 levels and as well as reduction in FGFR1 and FGFR4 levels (**Figure 1E and Supplementary Figure S1B**). As has been previously reported the PNT1A-HIP1 cell-line was characterised by significantly higher levels of FGFR4 (**Supplementary Figure S1C**). By contrast to our validation of STAT3 phosphorylation by Western blotting, it was not possible to detect increased levels of p-AMPK1α (T174), p-Akt (S473) or p-PLC-γ1 when we attempted to validate these by blotting (**Supplementary Figure S1D-F**). STAT3 has been associated with the transformation of an immortalized prostate epithelial cell line (14) to contribute to castration-resistance and the inhibition of apoptosis (15).

**Figure 1.**
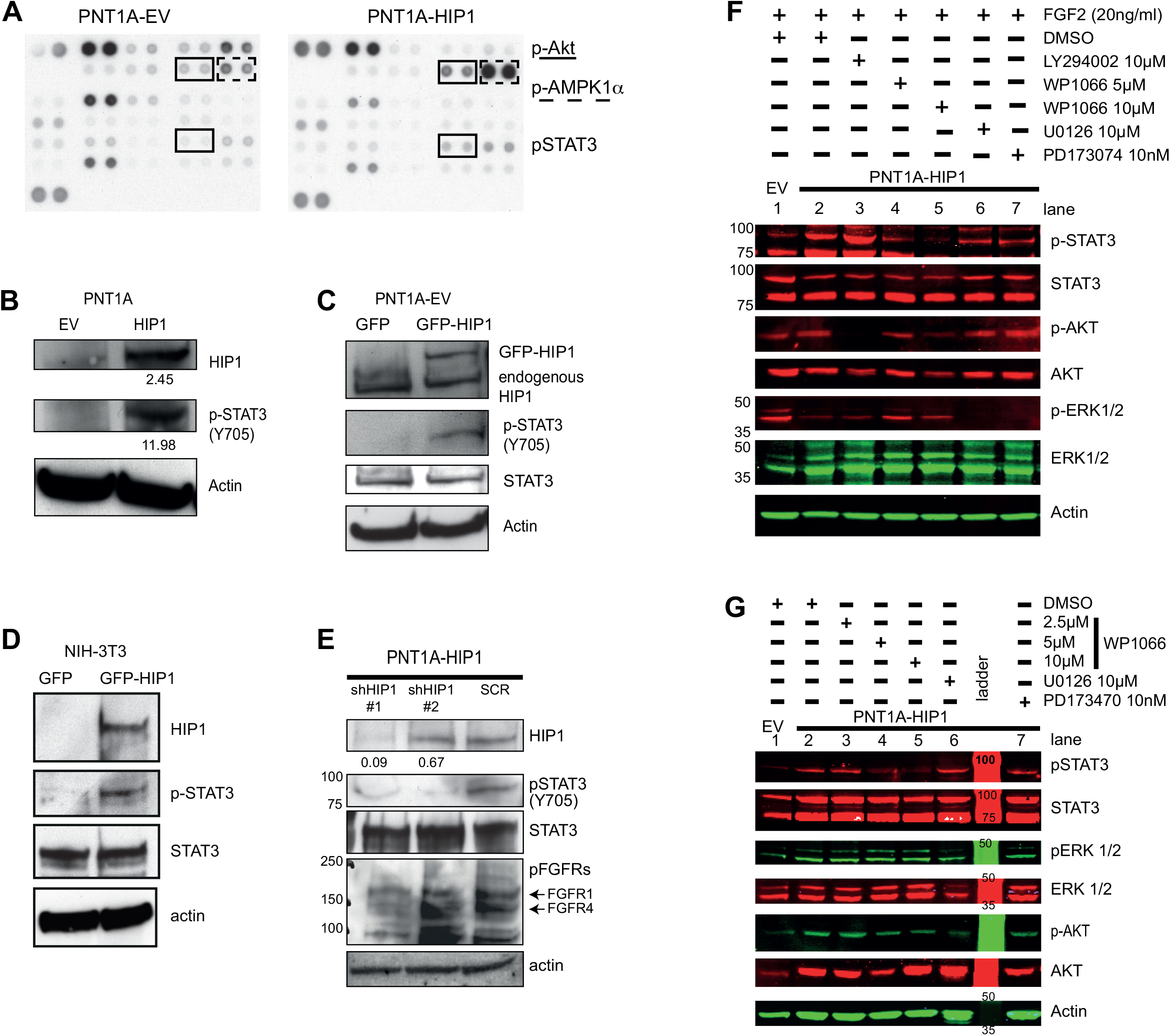
Increased STAT3 signalling with HIP1 overexpression occurs downstream of FGFR4 signalling and can be blocked with WP1066. **(A)** To identify downstream signalling pathways activated by HIP1 overexpression membrane A of the Human phospho-kinase array (ARY003) used to de-termine the relative levels of phosphorylated proteins of 46 specific kinases and substrates. Lysates were prepared from PNT1A-empty vector (EV) and PNT1A-HIP1 cell lines grown in full serum. The intensity of array spots in duplicate were analysed with ImageQUANT™ software. **(B)** Immunoblot of PNT1a-EV and PNT1a-HIP1 lysates for HIP1, p-STAT3(Y705) and actin. Relative levels of phosphorylation of STAT3 in PNT1a-HIP1 normalised to the control cell line after correction for equal protein loading control is displayed below blots. **(C)** Immunoblot of HIP1, p-STAT3(Y705) upon transient ectopic expression of GFP-HIP1 in the PNT1a-EV control for 48 hours. **(D)** Immunoblots of HIP1 and pSTAT3 in NIH3T3 cells transiently overexpressing HIP1. **(E)** Immunoblots of HIP1, p-STAT3(Y705), and p-FGFR following stable knockdown of HIP1 in PNT1a-HIP1 using shRNA compared to scrambled control. Fold changes indicated were normalised to actin. **(F)** WP1066 pre-treatment blocked STAT3 phosphorylation upon FGF2 stimulation in HIP1 overexpressing cells. PNT1A-HIP1 and PNT1A-EV were serum starved for 24 hours, pre-treated with DMSO or kinase inhibitors for one hour and stimulated with FGF2 (20ng/ml). Comparison of treatments with PI3K inhibitor (LY294002), Jak2 inhibitor (WP1066), MEK1/2 inhibitor (U0126), FGFR phosphorylation inhibitor (PD173074). **(G)** Drug treatment of PNT1A cells cultured in full serum conditions. PNT1A-EV and PNT1A-HIP1 cells cultured in full serum were treated with JAK2 kinase inhibitor (WP1066), MEK1/2 kinase inhibitor (U1026) and FGFR phosphorylation inhibitor (PD173074).

### Increased STAT3 phosphorylation in PNT1A-HIP1 is principally Jak2-mediated

Given the importance of FGFR signalling to the PNT1A-HIP1 cell-line we then investigated the relationship between FGF2-stimulated signalling and STAT3 phosphorylation in these cells. Serum starved PNT1A-HIP1 cells were pre-treated for one hour with inhibitors of PI3-kinase (LY294002), MEK1/2 (U0126), FGFR (PD173074), JAK2 (WP1066) or vehicle (DMSO). FGF2 was then applied for 30 minutes. In the vehicle control condition this led to an increase in STAT3 phosphorylation in the the PNT1A-HIP1 cells compared to PNT1A-EV (**Lanes 1 and 2; Figure 1E**). LY294002 blocked AKT phosphorylation (L**ane 3; Figure 1F**) without affecting p-STAT3 levels; WP1066 blocked STAT3 phosphorylation at both 5 and 10 μM concentrations (**Lanes 4 and 5; Figure 1F**); and U1026 and PD173074 blocked ERK1/2 phosphorylation whilst only partially affecting STAT3 phosphorylation (**Lanes 6 and 7; Figure 1F**).

We also investigated the role of JAK2-mediated STAT3 signalling in full serum conditions to account for the possibility that other mitogens might promote STAT3 phosphorylation. PNT1A-HIP1 and EV cells were treated with the same inhibitors for 24 hours and whole cell lysates were blotted for phosphorylated STAT3, ERK1/2 and AKT. WP1066 treatment blocked STAT3 phosphorylation at 5 and 10 μM concentrations (**Lanes 4 and 5; Figure 1G**) without affecting AKT and ERK1/2 phosphorylation levels. The ERK1/2 inhibitor U1026, and the FGFR phosphorylation inhibitor PD173074 did not affect STAT3 phosphorylation under steady state conditions (**Lanes 6 and 7; Figure 1G**).

To test whether FGF treatment increased the nuclear pool of STAT3, we quantified nuclear STAT3 levels using an imaging cytometer. PNT1A-EV and PNT1A-HIP1 cells were immuno-stained following 24 hours of serum starvation and FGF2 treatment for 30 minutes. FGF2 treatment of PNT1A-HIP1 significantly increased the nuclear pool of STAT3 whilst no significant difference in the nuclear pool of STAT3 was seen in the PNT1A-EV with FGF2 treatment when quantified relative to the surface area of nuclei (**Figure 2A**) and also when evaluated as the percentage of cells with nuclear staining for STAT3 (**Figure 2B**). Culturing PNT1A-EV cells in full serum significantly increased nuclear STAT3 as assessed by both parameters however the increase was greater for PNT1A-HIP1 cells cultured in the same conditions (**Figure 2A and 2B**).

**Figure 2.**
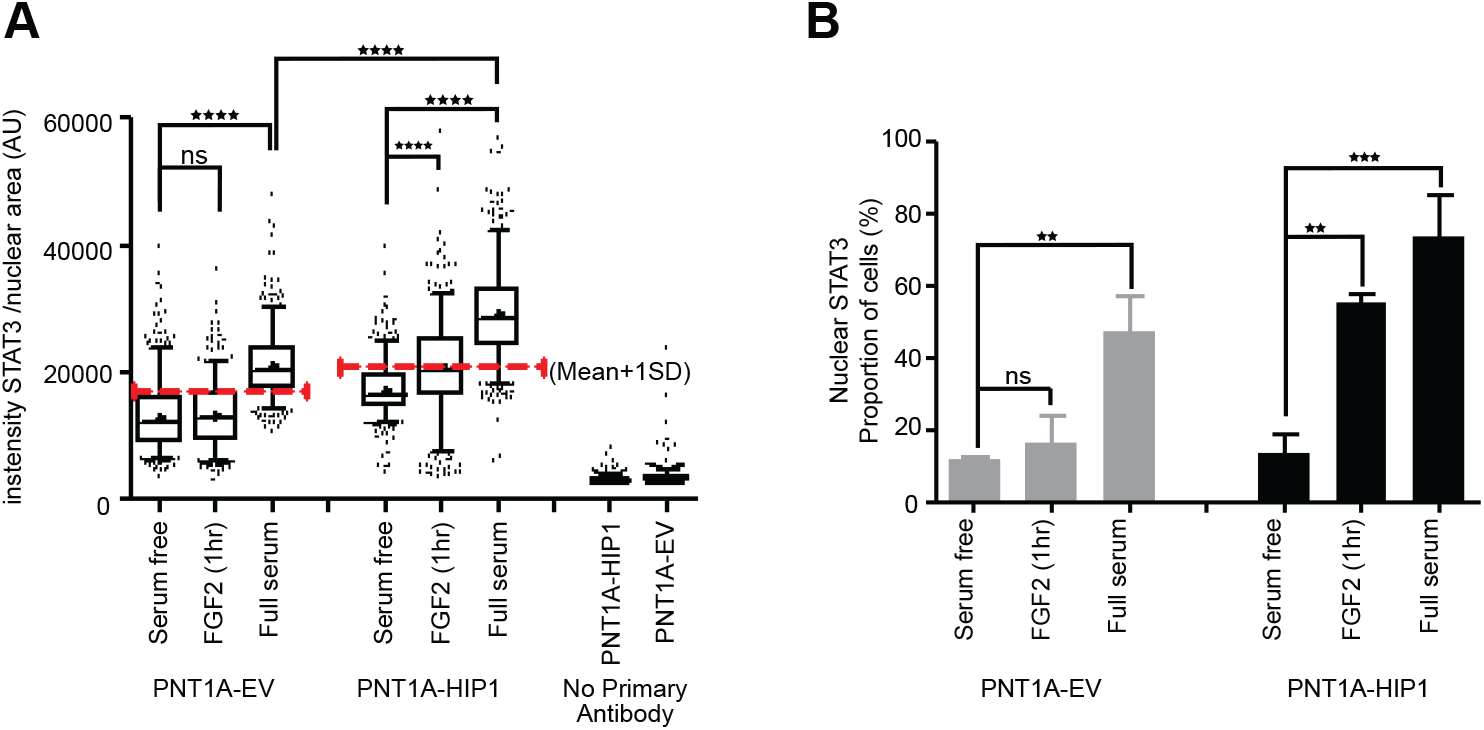
Mechanism and phenotypic effects of STAT3 phosphorylation in PNT1A-HIP1 cells. **(A)** Box plot showing median, 95% CI and mean ‘+’ of GFP-labelled nuclear STAT3 protein quantified using laser scanning cytometry; % of FGF2-treated and full serum culture cells with nuclear STAT3 > (mean nuclear STAT3 +1SD) in the control condition. Mean nuclear STAT3 levels assessed by laser scanning cytometry (iCYS). ****p<0.0001, n=500, one-way ANOVA was used to compare group. **(B)** Percentage of cells with mean nuclear STAT3 larger than mean + 1SD in respective serum free control. Data represents mean ± SEM, ***p<0.001, Students t-test, N=3 replicate experiments.

### STAT3 signalling is critical for the phenotypic effects of HIP1 overexpression

To test whether STAT3 mediated oncogenic transformation of PNT1A cells overexpressing HIP1, we inhibited STAT3 function by knocking down STAT3, as well as through the application of WP1066. Transient STAT3 knockdowns in PNT1A-HIP1 cells significantly reduced soft agar colony formation (**Figure 3A**) as did acute WP1066 at a 5μM dose (**Figure 3B**), suggesting a role for STAT3 signalling in the transformation phenotype seen in PNT1A-HIP1 cells.

**Figure 3.**
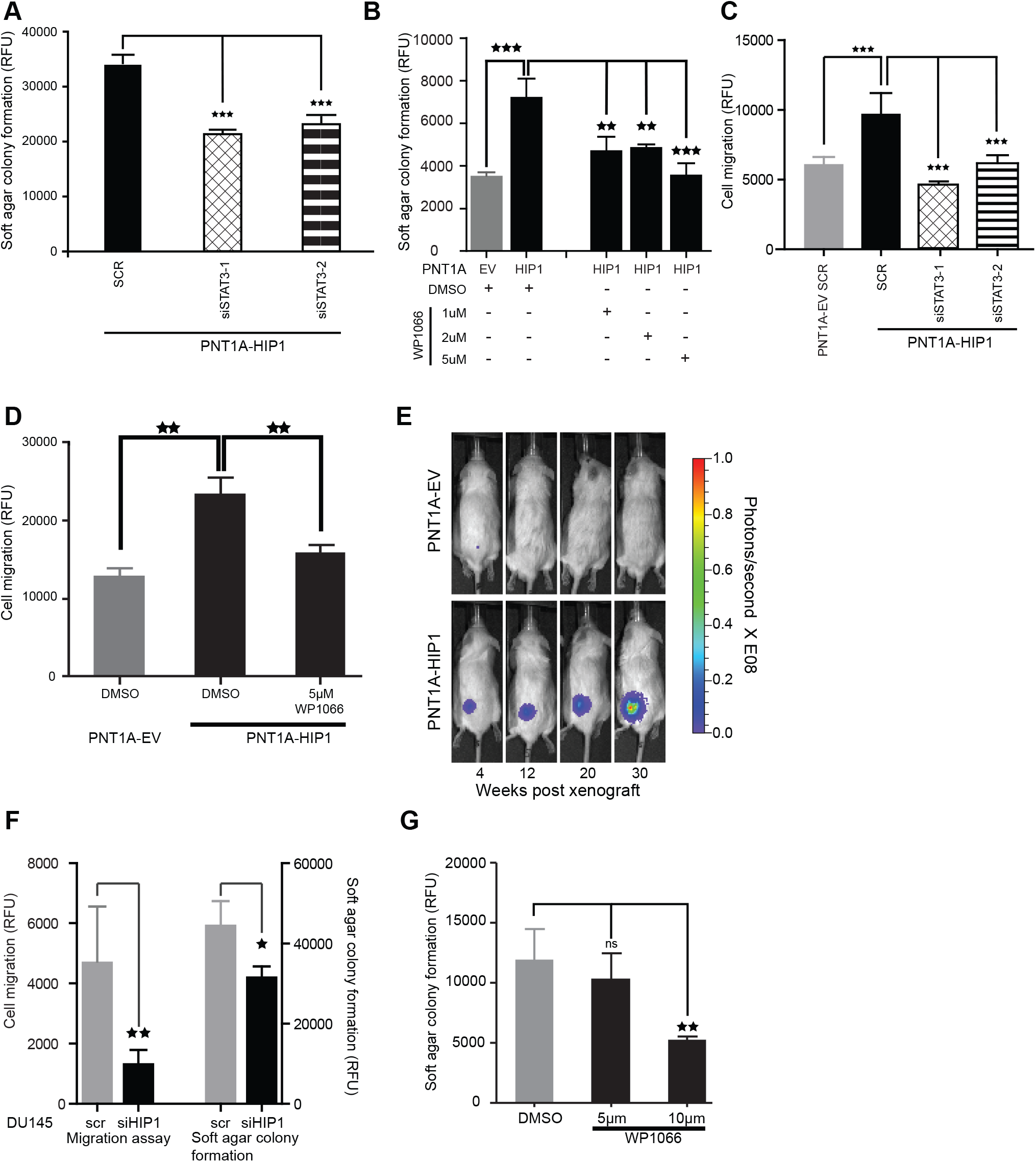
STAT3 signalling is critical to the phenotypic effects of HIP1 overexpression. **(A)** Soft agar colony formation assay for PNT1A-HIP1 cells following STAT3 knockdown. Data represents mean ± SD from 6 replicates in 3 independent experiments; ***p<0.001; Student’s t-test. Soft agar colony formation assay for PNT1A-HIP1 cells following treatment with STAT3 inhibitor WP1066. Data represents mean ± SD from 3 replicates in 3 independent experiments; ***p<0.001; **p<0.01; Student’s t-test. **(C)** Cell migration assay for PNT1A-HIP1 cells following STAT3 knockdown. Data represents mean ± SD, from 4 replicates in 3 independent experiments; **p<0.01; Student’s t-test. **(D)** Cell migration assay for PNT1A-HIP1 cells following treatment with STAT3 inhibitor WP1066. Data represents mean ± SD from 4 replicates in 3 independent experiments; **p<0.01; Student’s t-test. **(E)** *In vivo* tumour formation with PNT1A-HIP1 cells. Representative bioluminescent images of a mouse with PNT1a-EV (top) or PNT1a-HIP1 (bottom) imaged over 30 weeks post-xenograft. Scale represents bioluminescence in photons/sec. **(F)** Cell migration (left) and soft agar colony formation (right) was assessed 72 h post-transfection using Cellbiolabs^TM^ cell transformation assays of DU145 cells transfected with scrambled siRNA or pooled siRNA against HIP1. Colony formation was assessed 7 days following transfection, and is reflected by RFUs. Data represents mean ± SD from 3 replicates in 3 independent experiments; *p<0.05; **p<0.01; Student’s t-test. **(G)** Quantification of colony formation in soft agar in DU145cells treated with WP1066. Data represents mean ± SD from 4 replicates in 3 independent experiments; **p<0.01; Student’s t-test.

Next we investigated the effect of STAT3 inhibition on cell migration. PNT1A-HIP1 cells showed significantly greater cell migration compared with EV cells (**Figures 3C and 3D**). Both transient STAT3 knockdowns as well as WP1066 treatment at a 5μM dose significantly reduced PNT1A-HIP1 cell migration (**Figures 3C and 3D**).

The definitive test of cell transformation is the ability of xenografted cells to form tumours. To evaluate the tumorigenic potential of HIP1 overexpression we xenografted PNT1A-EV and PNT1A-HIP1 cell-lines stably expressing a luciferase reporter subcutaneously into mice. Tumour growth, assessed every four weeks by imaging, showed increasing bioluminescence of PNT1A-HIP1 xenografts, while PNT1A-EV xenografts failed to form tumours (**Figure 3E and Figure S2A**). Thirteen out of twenty PNT1A-HIP1 xenografts formed palpable tumours 30 weeks post-xenograft as exemplified in **Figure S2B**.

Next we extended our study to an additional prostate cancer cell-line, DU145, which has constitutive STAT3 activation and high HIP1 expression (6, 16). Upon knocking down HIP1 in DU145, we saw a significant reduction in soft agar colony formation and cell migration (**Figure 3F**). HIP1 knockdown also resulted in a reduction of p-STAT3 and a concomitant reduction in phospho-FGFR levels (**Figure S2C**). Inhibition of STAT3 signalling using 10μM WP1066 significantly reduced colony formation on soft agar further supporting the importance of an interplay between HIP1 and STAT3 in sustaining cellular phenotypes associated with tumorigenic potential (**Figure 3G**).

### GDF15 secretion reflects HIP1-mediated cellular transformation

To identify a conserved set of pathways that are affected by HIP1 overexpression we analysed gene expressing changes using gene expression microarrays. We undertook expression profiling on six replicates of PNT1A-HIP1, PNT1A-EV, LNCaP-HIP1, and LNCaP-EV. HIP1 overexpression was validated in all samples by rtPCR prior to expression profiling. Based on hierarchical clustering, five replicates of PNT1A-HIP1, PNT1A-EV, LNCaP-HIP1, and LNCaP-EV were used for further analysis. Using a p-value cut-off of <0.01 we identified 3,834 genes that were differentially expressed (DEG) in the PNT1A-HIP1 compared with the PNT1A-EV and 362 genes that were differentially expressed in the LNCaP-HIP1 cell line compared with the LNCaP-EV (**Figure 4A**). Of the differentially expressed genes we identified 112 genes that were overlapping among the PNT1A-HIP and the LNCaP-HIP1 datasets. Network analysis of the DEGs in these conditions highlighted STAT3 as a candidate transcriptional regulator associated with HIP1 overexpression in both LNCaP and PNT1A within regulatory networks including HIF1A in both cases (**Supplementary Figure 3**). Amongst the significantly downregulated genes with HIP1 overexpression in both lines were a significant number of genes associated with the innate immune/type 1 interferon response. By contrast amongst the most overexpressed genes in both cell-lines, GDF15, a TGF-beta superfamily receptor ligand, was the only common gene in the LNCaP and PNT1A overexpressing lines (**Supplementary Figure S4A**). GDF15 has been shown to be involved in prostate cancer progression, metastasis, and drug resistance to (17, 18). The overexpression of GDF15 was validated by rtPCR in the PNT1A cell line (**Figure 4B**). Further, stable HIP1 knockdown resulted in significantly lower GDF15 mRNA levels suggesting that GDF15 expression levels are dependent on HIP1 (**Figure 4B**). Western blot analysis of GDF15 protein expression in PNT1A-HIP1 cell lysates failed to identify GDF15 expression (**Supplementary Figure S4B**). However, since GDF15 is a secreted protein that undergoes significant processing in the Golgi network, we then quantified secreted GDF15 using an ELISA assay. A significant increase in GDF15 was observed in conditioned media (CM) derived from PNT1A-HIP1 compared to PNT1A-EV (**Figure 4C**). Furthermore, there was a significant reduction of secreted GDF15 in CM upon transient STAT3 knockdown or stable HIP1 knockdown in PNT1A-HIP1 **(Figure 4C**). The equivalent effect of HIP1 and STAT3 knockdown on the levels of secreted GDF15 highlight the contribution to GDF15 regulation. We also found that WP1066 treatment significantly reduced GDF15 secretion in the PNT1A-HIP1 cell lines (**Figure 4D**).

**Figure 4.**
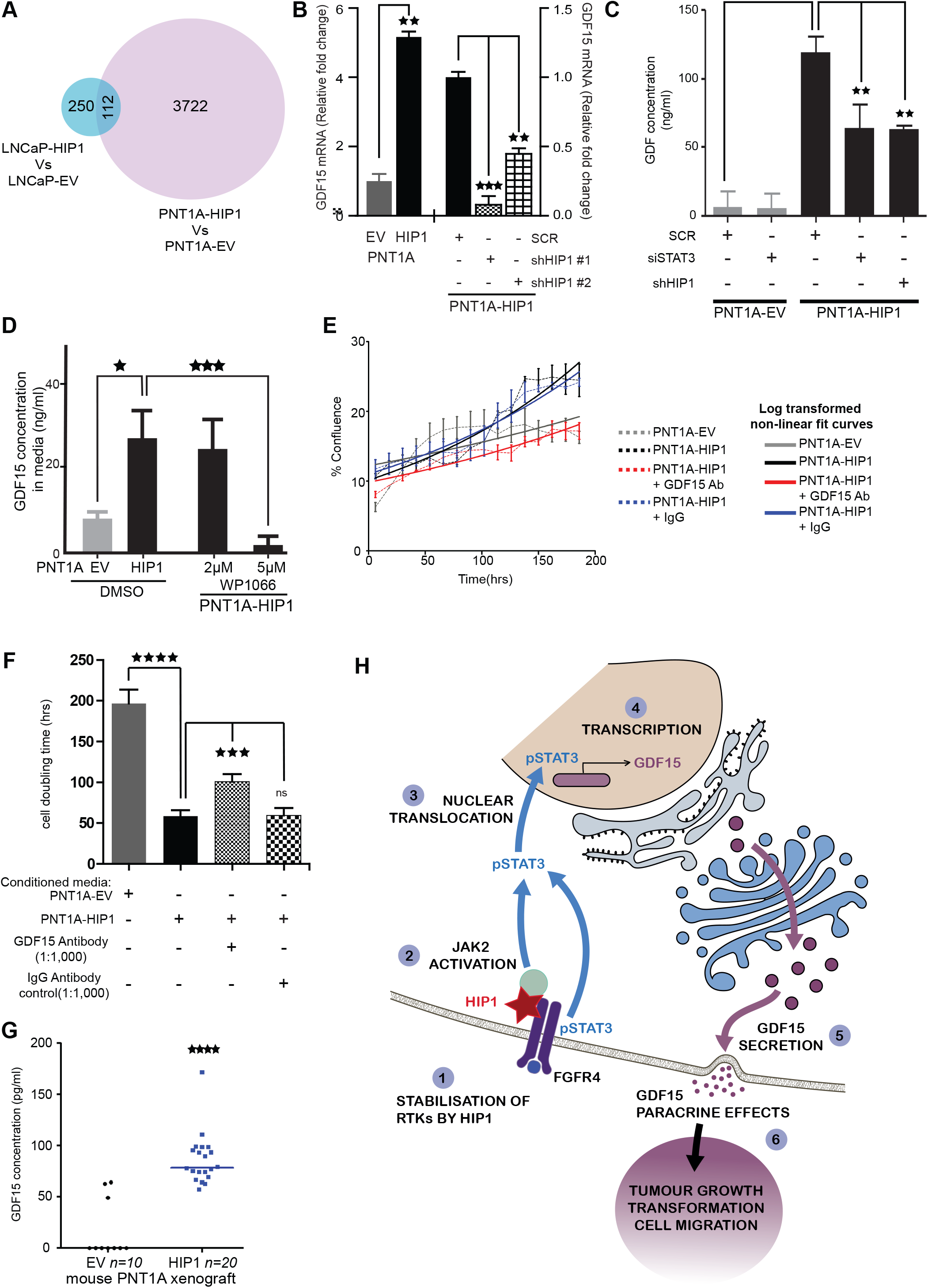
Assessment of HIP1 and serum GDF15 as biomarkers of prostate cancer. **(A)** Venn diagram showing differentially expressed genes in HIP1 overexpressing PNT1A and LNCaP cell lines. **(B)** Quantification of GDF15 mRNA in PNT1A-HIP1 cells compared to control (left) and upon HIP1 knockdown with shRNA (right). Fold changes were assessed by comparing PNT1A-shHIP1 with control. The experiment was performed 3 times; **p<0.01; ***p<0.001; Student’s t-test. **(C)** Quantification of secreted GDF15 in conditioned media (CM) from PNT1a-EV and PNT1a-HIP1 analysed in triplicate in 3 independent experiments. One-way ANOVA was used to assess the differences in GDF15 secretion between the various groups; **p<0.01. **(D)** Quantification of secreted GDF15 in CM from PNT1a-EV and PNT1a-HIP1 following 48 hr treatment with DMSO or 2μM, and 5μM STAT3 inhibitor WP1066. Data represents mean ±SD of experimental triplicates in three independent experiments. One-way ANOVA was used to assess the differences in GDF15 secretion between the various groups; *p<0.05; ***p<0.001. **(E)** Parental LNCaP cell lines were seeded at equal concentration in a 24-well plate in duplicate(n=2). 24 hours later, CM (day 5) from PNT1A-EV or PNT1A-HIP1 was added and the cells were cultured; growth of LNCaP was measured as the degree of confluence per well area. Data represents mean*±*SD of two wells (n=2). CM from PNT1a-HIP1 was pre-treated with a GDF15 antibody or IgG control antibodies for 1h prior to addition to the cultures. Data was analysed using 4-parameter log non-linear regression curve fitting. **(F)** Doubling times for PNT1A-HIP1 cells grown in conditioned media treated with anti-GDF-15 antibodies or IgG control. Data represents mean ± SEM of 4 independent experiments; ***p<0.001; ****p<0.0001. **(G)** GDF15 was detectable in mice sera bearing PNT1A-HIP1 xenografts but not in the EV-xenograft mice sera. About 100*μ*L of serum was collected from the mice from the jugular vein 26 weeks post-xenograft of PNT1A-HIP1 (n=20) and PNT1A-EV (n=10). Scatter plots representing each mouse serum GDF15 concentrationand median serum GDF15 concentration (horizontal line) has been shown. Mann-Whitney U test was used to compare the data; ****p<*<*0.0001. **(H)** Schematic diagram illustrating the mechanism of HIP1 in cancer progression via STAT3 signalling and GDF15 secretion.

### Paracrine effects of GDF15 may contribute to the role of HIP1 in cancer progression

Here we have demonstrated HIP1 overexpression to be linked with increased GDF15 secretion. To test whether factors secreted by the PNT1A cell-line may have paracrine effects, we treated the LNCaP cell-line cultured in the absence of androgens with conditioned media from PNT1A-HIP1 and PNT1A-EV. As a control, GDF15 was depleted from the conditioned media with anti-GDF15 antibodies or control IgG antibodies prior to the media being applied to LNCaP cells. LNCaP cells cultured with conditioned media from PNT1A-HIP1 had increased growth rates (**Figure 4E**) and significantly lower doubling times compared to those grown with conditioned media from PNT1A-EV (**Figure 4F**). Moreover, depleting GDF15 from conditioned media from PNT1A-HIP1 cells significantly abrogated the enhanced growth induced in the LNCaP cell-line upon exposure to these media (**Figures 4E and 4F**). By contrast mock depletion with control IgG antibody did not affect the growth advantage induced by conditioned media harvested from PNT1A-HIP1 cells.

### GDF15 is a biomarker of HIP1-driven *in vivo* cell transformation

As shown previously, PNT1A-HIP1 xenografts formed tumours in mice, while PNT1A-EV xenografts failed to do so (**Figure 3E**). To test whether GDF15 secretion was also detectable with cellular transformation induced by HIP1 overexpression *in vivo*, we assessed serum GDF15 concentration in mice sera following establishment of tumours at 30 weeks. In accord with our *in vitro* findings, serum GDF15 concentration in PNT1A-HIP1 xenografted mice was significantly higher compared to the age-matched PNT1A-EV xenografts. GDF15 was undetectable in the majority of PNT1A-EV xenografted mice (**Figure 4G**).

## Discussion

HIP1 has previously been reported to stabilize receptor tyrosine kinases (RTKs) including EGFR and FGFR4 by promoting their recycling through a clathrin-mediated endocytic pathway (9, 12). Overexpression of HIP1 in an NIH-3T3 cell-line leads to increased tumorigenicity of this line by sustaining kinase signaling cascades downstream of RTKs even under conditions of serum deprivation(12). HIP1 is also known to be overexpressed in tumour tissue from prostate cancer, colorectal cancer and a number of other cancer types in and this elevated expression associates with poor patient outcome/disease progression(6). In this study we have further interrogated the downstream pathways affected by HIP1 overexpression in a prostate cell-line, PNT1a, which are immortalized prostatic epithelial cells. This overexpression model is one in which FGFR4 has previously been found to be more highly expressed and this has been associated with a physical interaction between FGFR4 and HIP1(9). Through the use of a phospho14-kinase antibody array we have found that phosphorylated STAT3 is elevated in this model and we have gone to show that its phosphorylation correlates with its increased nuclear localization (**Figure 1 and Figure 2**). These characteristics were reversed by knocking down HIP1 and could also be induced by transient HIP1 overexpression the PNT1A-EV line and also the NIH-3T3 cell-line (**Figure 1**). Furthermore, phenotypic features of HIP1 overexpression including colony formation on soft agar and increased cell migration could all be reversed by treatment with a STAT3 inhibitor and by STAT3 knockdown (**Figure 3**). Knocking down STAT3 or HIP1 in a cancer cell-line, DU145, also reduced the migratory capacity of this line. Collectively these data indicate that STAT3 activation is an important mediator of cellular transformation driven by HIP1 overexpression.

FGFR4 has been reported to be a direct binder of STAT3 and inducer of STAT3 phosphorylation. We also know that activating mutations in FGFR4 associate with poor prognosis disease, particularly of arginine-388, and functionally this stabilizes the FGFR4-STAT3 interaction and enhances STAT3 phosphorylation(19). Since HIP1 overexpression also enhances FGFR4 expression and has previously been potentiate FGFR signaling(9) our finding that this also induces STAT3 phosphorylation ties these hitherto distinct strands of research together.

This has potentially important implications for downstream signaling. Our microarray data from both PNT1a and LNCaP cell-lines overexpressing HIP1 show that type 1 interferon response/innate immune genes are downregulated compared to empty vector control cells. STAT3 is known to have repressive effect on the transcription factor complex that drives the expression of type 1 interferon response genes, a complex known as ISGF3(20, 21). ISGF3 is a heterotrimer of STAT1, STAT2 and IRF9 and its activation can in cancer cells reflect the presence of unresolved DNA damage in the form of elevated cytosolic DNA and also aberrant transcription of retro-transposons and from endogenous retroviral insertion sites(22, 23). Inducing the activation of this complex is an increasingly therapeutic strategy for the improving the response of cancer patients to therapy and has been linked to the use of epigenetic inhibitors, such as DNMT1 and HDAC-targeted drugs, as well agonists of the nucleic acid sensing pattern recognition receptors such as STING(24, 25). One implication of our work therefore is that HIP1 overexpression may also support the immune evasive properties of cancer cells. This hypothesis is further supported by the finding that a germline mutation in FGFR4 (rs351855) which enhances STAT3 signalling was recently found to promote immune evasion in transgenic knock-in models of lung and breast cancers(26). Given that FGFR4 interacts with HIP1 and we have shown HIP1 overexpression enhances STAT3 signalling it will be important to evaluate the immunological effects in syngeneic or transgenic cancer models in the future.

In addition, we have found that a TGF-beta superfamily ligand, GDF15/MIC-1, with established immunosuppressive properties and associated with tumour cachexia, is significantly elevated in HIP1 overexpressing cells and this depends on STAT3 activation (**Figure 4**). It too has been found to restrict the immune response in immune-competent cancer models for example by inhibiting T cell infiltration in glioblastoma(27, 28). Recently its receptor was identified in brain, GFRAL, and in this context is being pursued as a target for the treatment of obesity(29). It functions as an active signaling complex in association with RET, its coreceptor, upon GDF15 binding(30). The contribution of this complex to cancer development is currently largely unexplored although RET on its own is an established therapeutic target for a number of cancer types including prostate cancer(31, 32). A second important avenue for future study will be the relevance of this complex to prostate cancer progression and the impact of HIP1 expression on its trafficking and activity. All of this work will require multiparametric analysis in clinical samples and the ability to do this for protein markers in tissue at a molecular level is progressing rapidly through the development of mass imaging cytometry and other technologies. FGFR4 itself is by contrast a much more established receptor target and is well-characterized in many cancers as a driver of treatment resistance. In colorectal cancer, another cancer type in which HIP1 overexpression is a poor prognosis biomarker, FGFR4 is known to support STAT3 activation with the concomitant overexpression of anti-apoptotic genes such as CFLAR which encodes c-FLIP(33). It will therefore be interesting to determine what the repertoire of genes and pathways is, which functionally associate with both HIP1 overexpression and STAT3 activation to drive disease progression in other cancer types. It is likely that some will be cancer type-specific and some will be cancer cell autonomous hallmarks of these factors.

## Conclusions

In conclusion, we have shown for the first time that the oncogenic potential of HIP1 overexpression is mediated by STAT3 activation acting in part through the increased secretion of GDF15 (**Figure 4H**). Experiments demonstrated that HIP1 overexpression increased STAT3 signalling in response to FGF2 receptor activation and increased GDF15 transcription. The increase in GDF15 protein secretion was dependent on HIP1 and STAT3 expression and was shown to have paracrine growth-promoting effects. The HIP1-STAT3-GDF15 axis provides new insight into the mechanisms of prostate cancer development.

## Abbreviations

HIP1: Huntingtin-interacting protein 1
CaP: prostate cancer
STAT3: Signal transducer and activator of transcription 3
kDa: kilo Dalton
FGF: fibroblast growth factor
FGFR: fibroblast growth factor receptor
GDF15: Growth Differentiation Factor 15
AR: androgen receptor
GFP: green fluorescent protein
HRP: horse radish peroxidase
qRT-PCR: quantitative real time polymerase chain reaction
GAPDH: Glyceraldehyde 3-phosphate dehydrogenase
EV: empty vector
shRNA: short hairpin ribonucleic acid
RPMI: Roswell Park Memorial Institute
FCS: foetal calf serum
mRNA: messenger ribonucleic acid
siRNA: Small interfering ribonucleic acid
PMSF: Phenylmethanesulfonyl fluorid
DTT: Dithiothreitol
EDTA: Ethylenedinitrilotetraacetic acid
SDS-PAGE: sodium dodecyl sulfate-polyacrylamide gel electrophoresis
*PVDF*: *Polyvinylidene difluoride*
RFU: relative fluorescence unit
AMPK1α: Protein kinase AMP-activated catalytic subunit alpha 1
PLC-γ1: Phospholipase C-γ1
MEK1: Mitogen-activated protein kinase kinase 1
JAK2: Janus kinase 2
DMSO: Dimethyl sulfoxide
ERK1/2: extracellular signal-regulated protein kinase ½
DEG: differentially expressed gene
TGF-beta: transforming growth factor-beta
ELISA: enzyme-linked immunosorbent assay
IgG: Immunoglobulin G
RTKs: receptor tyrosine kinases
EGFR: epidermal growth factor receptor
ISGF3: Interferon stimulated gene factor 3
IRF9: Interferon regulatory factor 9
DNMT1: DNA (cytosine-5)-methyltransferase 1
HDAC: Histone deacetylase
STING: stimulator of interferon genes
GFRAL: Glial cell-derived neurotrophic factor Family Receptor Alpha Like
RET: rearranged during transfection
CFLAR: CASP8 and FADD-Like Apoptosis Regulator

## Declarations

### Ethics approval

All animal procedures were carried out in accordance with University of Cambridge and Cancer Research UK guidelines under UK Home Office project license 80/2301.

### Consent for publication

Nothing to declare as no patient material has been used.

## Availability of data and materials

The expression microarray data generated in this study have been deposited and can be accessed through the NCBI GEO Dataset repository (https://www.ncbi.nlm.nih.gov/geo/) through accession number GSE35211.

## Authors’ contributions

RBR performed the experiments, analysed the data and drafted the manuscript. AR-M designed and performed the *in vivo* experiments, analysed data and contributed to the manuscript preparation. HS provided technical support for the laboratory work and contributed to the manuscript preparation. LB and SR provided guidance on the image analysis undertaken within the Light Microscopy Core Facility run by SR within the CRUK Cambridge Institute. SM and RR provided supported the analysis and deposition of gene expression data through the Bioinformatics Core Facility within the CRUK Cambridge Research Institute. EE contributed to the writing of the manuscript and data interpretation. DEN and IGM supervised the project and were responsible for overseeing the manuscript.

## Acknowledgements

We thank the core facilities at the Cancer Research UK Cambridge Institute led by James Hadfield (Genomics), Matt Eldridge (Bioinformatics) and Allen Hazelhurst (BRU), as well as Jodi Miller (IHC), Ellie Pryor (BRU) and Jason Fox (BRU). We would like to thank Michael Ittmann for providing the PNT1A lines; Scott Lyons for the luciferase-YFP pCyL50 plasmid used to generate xenografts. None of the authors has any conflict of interests.

