## Supplementary legends and figures S1 to S4 for "HIP1 mediates oncogenic transformation and cancer progression through STAT3 signalling"

***Supplementary information***

**Supplementary Figure S1. *HIP1 overexpression results in STAT3 signalling.* (A)** Quantification of Human Phospho-kinase (R&D Systems, ARY003) array revealed for PNT1A-HIP1. PNT1A-HIP1 array blot was compared with the PNT1A-EV blot to determine fold changes of phospho-proteins. A cut-off fold change of>1.5 (red line) in the PNT1A-HIP1 cell line was used to obtain a phosphorylation profile relating to HIP1 overexpression. (**B**) Relative HIP1 mRNA expression was determined by real time PCR of PNT1A treated with HIP1 shRNA and control shRNA. Data represent mean ± SD of experimental triplicates (N=3) normalised to housekeeping genes. (**C-F**) Western blot analysis of cell lysates from PNT1A-HIP1 and control PNT1A-EV cell lines. Protein phosphorylation of known downstream effectors of FGFR signalling were tested by western blotting for (**D**) ERK1/2 and PLCγ1 and (**E**) AKT. (**F**) Validation of Human Phospho-kinase (R&D Systems) array by Western blotting of PNT1A-HIP1 and DU145 cell lysates in duplicates.

**Supplementary Figure S2*. Xenografting of PNT1A-HIP1 cells results in tumour growth.* (A)** To assess the formation/ growth of tumours following engraftment of PNT1A-HIP1 and PNT1A-empty vector, bioluminescent imaging (BLI) of the mice following IP injection of luciferin (150mg/kg) was conducted serially, every four weeks. Two-way ANOVA was used to assess the differences in bioluminescence between PNT1a-HIP1 and EV. The final reading was taken 30 weeks following xenografting. **p<0.01, ****p<0.0001. N=3 **(B)** Images showing tumour growth 30 weeks post-xenograft. **(C)** Western blot analysis of pSTAT3 in cell lysates from DU145 cells treated with siRNA targeting HIP1 or a scrambled control.

**Supplementary Figure S3. *Network analysis of differentially expressed genes in cell lines overexpressing HIP1.*** Network analysis of DEG lists was carried out using proprietary gene expression analysis software - GENEGO MetacoreTM. A visual representation of the two top-scored networks (p<0.01) obtained from analysis of **(A)** PNT1A-HIP1 vs EV comparison DEGs, **(B)** LNCaP-HIP1 vs EV DEGs, **(C)** DEGs common to LNCaP-HIP1 vs EV and PNT1A-HIP1 vs EV comparisons. The STAT3 network is highlighted with a green box.

**Supplementary Figure S4*. Identification of GFD15 overexpression in cell lines overexpressing HIP1.*** **(A)** Top 10 upregulated genes in LNCaP-HIP1 vs EV and PNT1A-HIP1 vs EV array comparison. DEG list for LNCaP-HIP1 vs EV and PNT1A-HIP1 vs EV array comparisons were computed using Bioconductor for R. A cut-off p-value of<0.01 was used to identify DEGs. DEG list was then sorted based on log fold change (FC) with overexpressed genes having positive FCs and downregulated gene having negative FCs. The top 10 overexpressed genes in each of the comparisons has been shown here with the log fold change, and p-value adjusted for multiple comparisons (adj.p-Val). **(B)** Western blots for GDF15 protein in PNT1A-HIP1 and PNT1A-EV cell lines. GDF15 western blots failed to detect GDF15 protein expression. PNT1A-HIP1and EV cells cultured in steady state were used to prepare lysates.

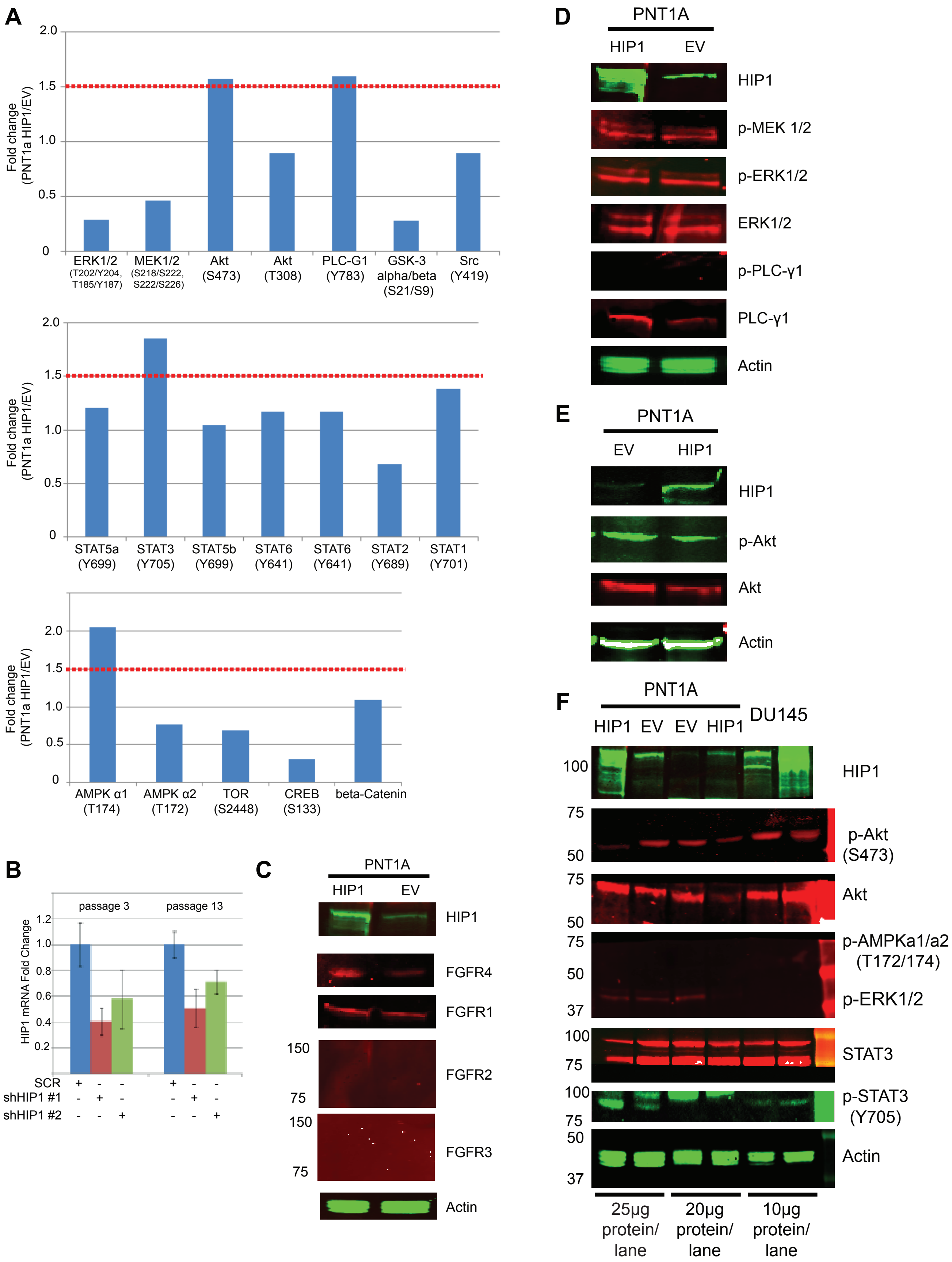


**Supplementary Figure 1**

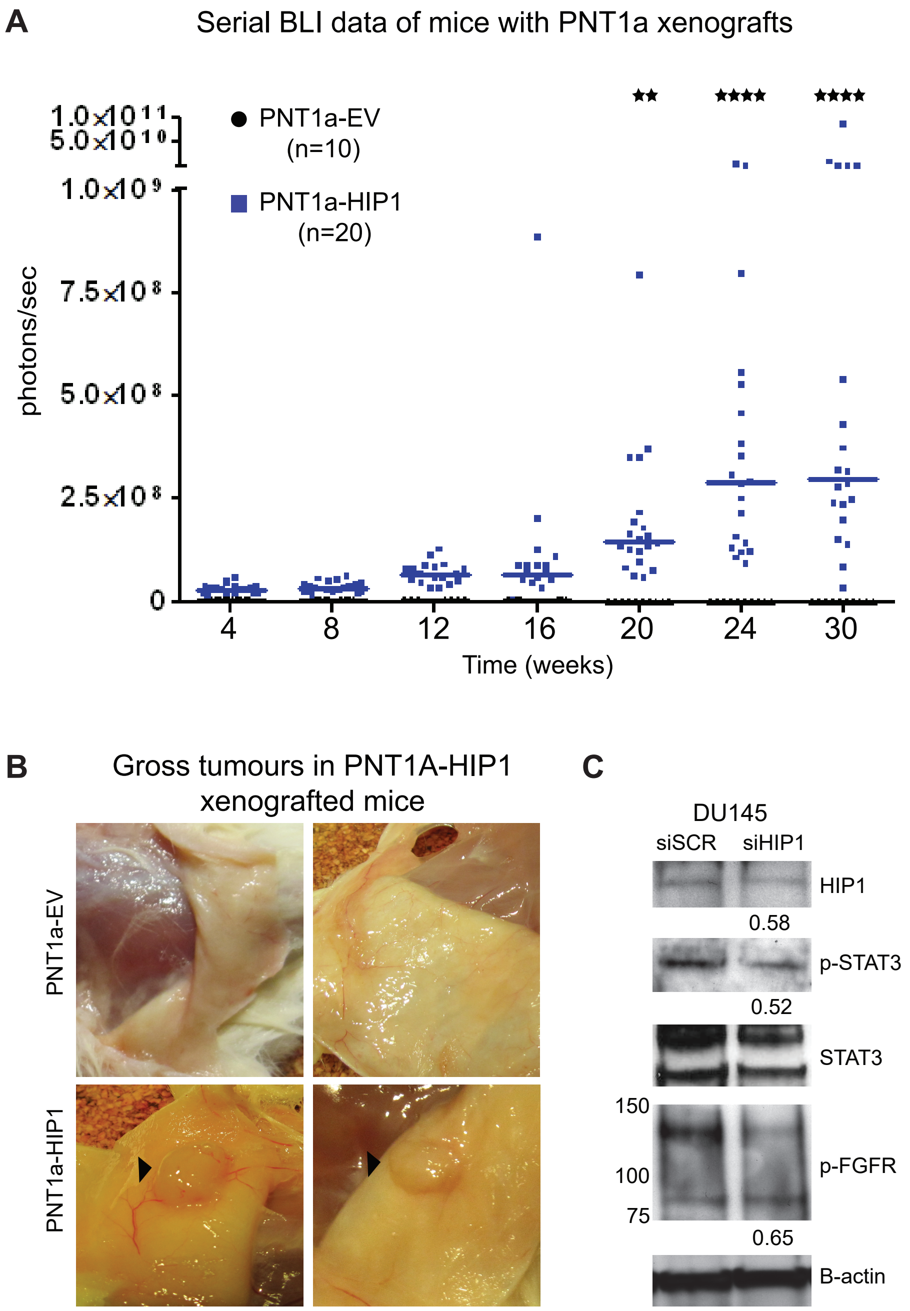


**Supplementary Figure 2**

### Top scored networks (p<0.01)

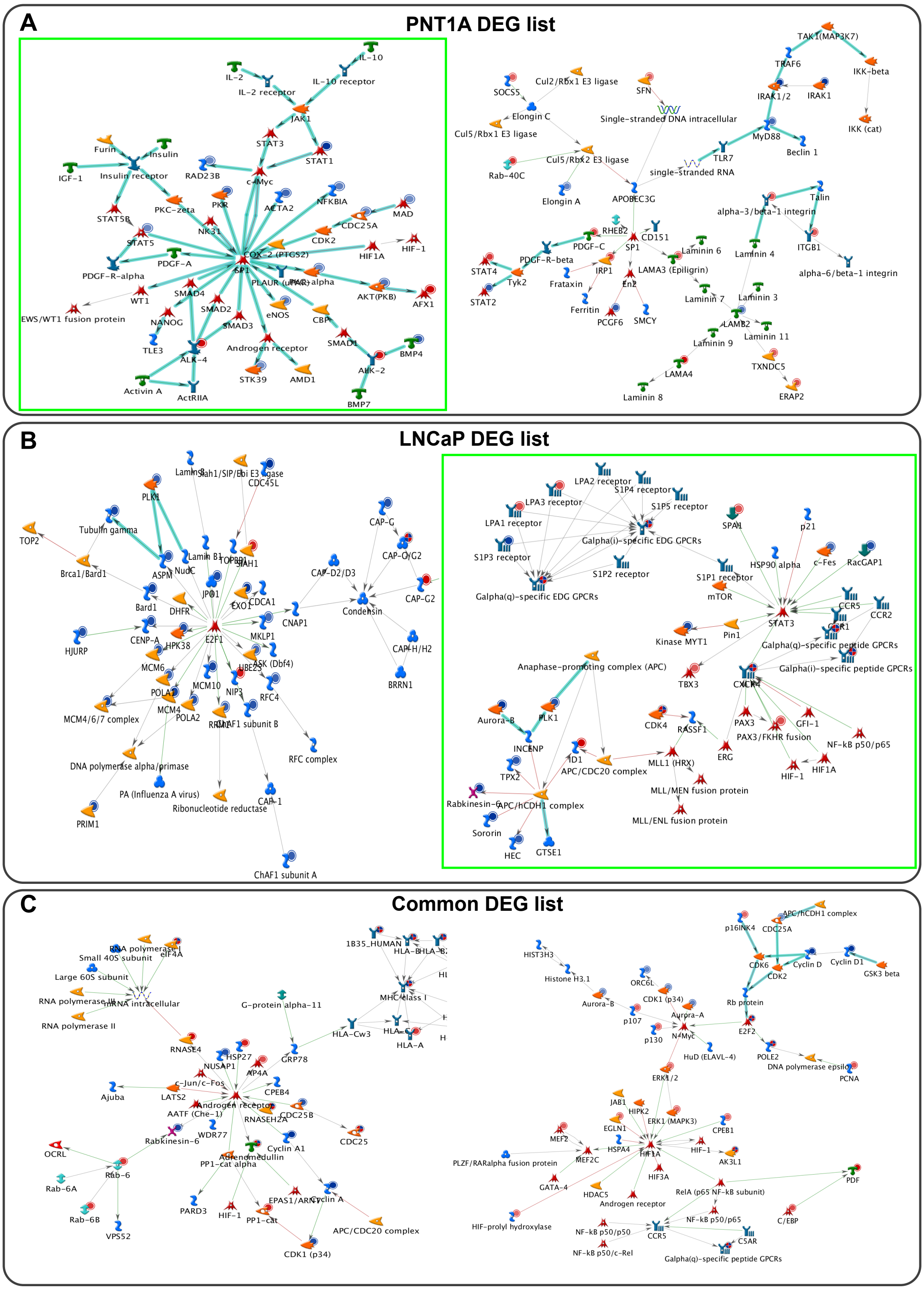


Supplementary Figure 3

**A**

LNCaP-HIP1 vs EV

| *Gene Symbol* | *logFC* | *adj. p-Val.* |
|---|---|---|
| NDUFA4L2 | 0.880 | 3.87E-05 |
| **GDF15** | 0.814 | 7.52E-06 |
| NDRG1 | 0.771 | 1.01E-05 |
| ENO2 | 0.743 | 8.40E-06 |
| DHRS3 | 0.701 | 4.05E-04 |
| BCHE | 0.647 | 3.84E-09 |
| CXCR4 | 0.638 | 7.95E-06 |
| P4HA1 | 0.620 | 7.66E-06 |
| NYPR1 | 0.619 | 1.59E-05 |

PNT1A-HIP1 vs EV

| *Gene Symbol* | *logFC* | *adj. p-Val.* |
|---|---|---|
| S100A9 | 3.698 | 1.72E-18 |
| RUNX3 | 3.616 | 5.10E-31 |
| SPOCK1 | 3.554 | 1.38E-27 |
| SAA1 | 2.634 | 7.42E-12 |
| NPY | 2.356 | 3.47E-26 |
| DDIT4 | 2.343 | 2.49E-13 |
| MDK | 2.175 | 1.95E-18 |
| FBLN2 | 2.161 | 6.50E-23 |
| **GDF15** | 2.117 | 6.23E-15 |

**B**

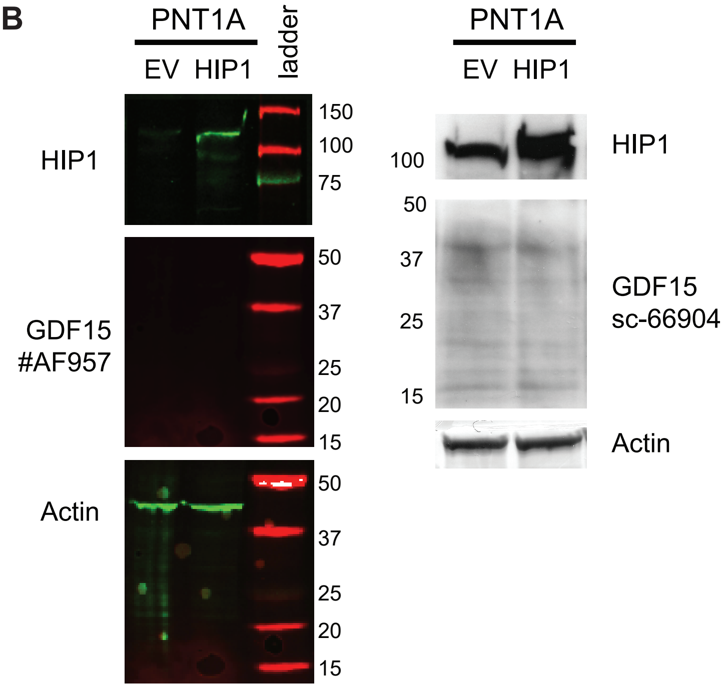


**Supplementary Figure 4**
